# Evolving Structure and Diversity of Human Narratives in the Journal of Stories in Science

**DOI:** 10.1101/2020.12.08.417071

**Authors:** Reilly Q. Mach, Jessica W. Tsai, Fanuel J. Muindi

## Abstract

Fundamentally, science is about people. The stories of the people behind the science are just as important as the results themselves, even though the results are often what get more attention. Despite the growth of databases storing such stories across diverse mediums, a detailed assessment of those databases is missing. To continue innovating science storytelling, we provide the first assessment of the structure and diversity of the narratives published in the Journal of Stories in Science. In this assessment, a total of 170 published stories authored by 158 authors between 2016 and 2020 are analyzed. Majority (67%) of the authors are women from North America and in the life sciences. The narratives most commonly feature authors from academia (e.g., 23% graduate students, 13% post-doctoral fellows and 21% professors). However, there is also a growing number of authors with PhDs that are working outside of academia (15%). Nearly a quarter (23%) of the database authors come from racial groups (African American, Latino, and Hispanic) that have been shown to be underrepresented in health-related sciences in the United States. Using the industry standard Flesch Reading Ease Score, we found that 74% of the stories included in the analysis fall in the target range of 50-70, which represents readability by students in grades 8-12. The analysis here provides the first deep look into one of the databases publishing diverse stories in science using a wide range of mediums. In summary, there is a need for more emphasis on both expanding and studying such databases given the continuous demand for these stories and their inclusion into K-12 curriculums.

## Introduction

At its core, science is about people. It is easy to get caught up in the results and publications, but there is always more than what is said in the data. Typical science storytelling involves researchers finding the best way to communicate their findings into a product that will be featured in a peer-reviewed scientific publication.^1^ Although this form of science writing is imperative, it fails to convey the humanity in science. The stories behind the science are just as, if not more, important than the results themselves.^2^ However, these stories are not being told. It is becoming increasingly important for scientists to be able to tell the human aspects behind their scientific work, rather than solely focusing on publishing their data.^3^ The emphasis on results can lead to the all too prevalent idea that scientists must follow a clear, linear path in order to be successful. This is simply not the case. Telling the human aspects behind the science helps diminish the idea that science is only for individuals with a defined set of qualities and accomplishments. Publishing these stories is an incredibly important aspect of science that must continue to gain recognition.

As scientists tell their stories, it is also imperative that their narratives are examined as they are stored in the growing number of online databases.^4^ A number of questions arise. For example, who are the authors? What is their career stage? Where are they located? What are their stories about? What is the reach and impact of their stories? How accessible are their stories? Are the stories increasing our understanding of what it means to be a human in science? What is the feedback from the readers? These are just some of the growing number of questions that need addressing. Despite the additional growth of databases storing such stories across diverse mediums, a detailed assessment of these stories is missing.

The *Journal of Stories in Science* is one such database that the Science Advocacy Institute (SAi) started in 2016 with the mission of publishing inspiring, thought-provoking, and human-centered stories in science from scientists around the world. With a current database of almost 200 stories, we provide the first formal assessment of the structure and diversity of the published stories thus far. Additionally, we also share insights into new areas of growth for both the journal and the field overall.

### Overview of the database

The journal typically publishes 3-4 new stories per month with an average length of 1000 words per publication. In this assessment, a total of 170 published stories authored by 158 authors (some authors published more than one story) between December 3, 2016 and June 14, 2020 are analyzed. Some of the authors’ stories focus heavily on their scientific work, while others write about what drew them into science in the first place, and expand on the progression of their careers in science thus far. The standard arc of most stories is typically the same: the exposition where the authors set the stage; the rising conflict where the authors share specific incidents or periods of time where they faced significant challenges; and the resolution where the authors discuss how they overcame the aforementioned challenges and setbacks that allowed them to move forward in their scientific journey. The published stories are tagged across several domains including author location, gender, employment sector, career type and also trainee status among others. The search feature on the site uses the restrictive AND logic function which makes it easier for a reader to quickly find the one or two stories they are looking for. In this assessment, a total of 170 published stories authored by 158 authors (some authors published more than one story) between December 3, 2016 and June 14, 2020 are analyzed.

### Structure and Diversity

As the number of published stories continues to grow, it is essential to track both the structure and diversity of the database in order to ensure that readers have access to a broad spectrum of authors’ stories. In order to study the developing structure of the database, we conducted a demographic analysis of 158 authors across four major categories: the primary author’s gender, location, field, and career stage at the time of writing. Overall, the author demographics are shown by Figure 1. We found that a majority of the authors are women from North America and in the life sciences (Figure 1A, 1B, 1C). It is important to note that the editorial team is made up of life scientists, which likely contributes to this bias. While the team tries to keep the database as diverse as possible, life scientists tend to be more attracted to the journal. Another interesting point comes from the gender breakdown, as shown in Figure 1A, with the majority of stories (67%) published by women. We also conclude that the database most commonly features authors from academia (e.g., 23% graduate students, 13% post-doctoral fellows and 21% professors) (Figure 1D). However, there is also a growing number of authors with PhDs that are working outside of academia (15%). It is worth mentioning that nearly a quarter (23%) of the database authors come from racial groups (African American, Latino, and Hispanic) that have been shown to be underrepresented in health-related sciences in the United States^5^ (Figure 1D).

**Figure 1:**
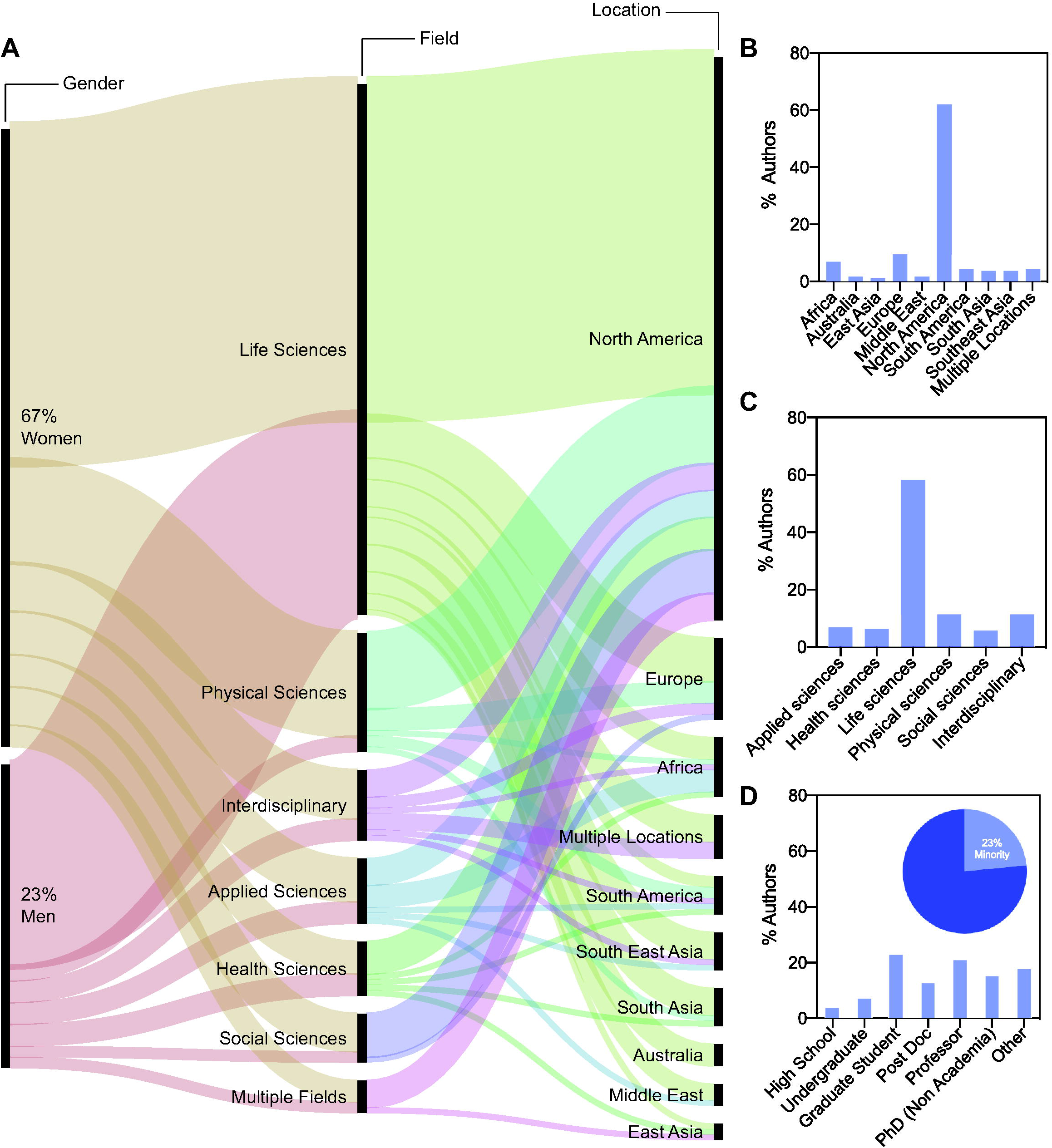
Author demographics. **(A)** An alluvial diagram generated by Rawgraphs.io graphically represents the changes across three key author demographics represented by the following vertical black lines: gender, field, and location. The width of each colored horizontal line is proportional to the number of authors within each cluster. **(B)** The percentage of authors across different locations. **(C)** The percentage of authors across different fields. **(D)** The percentage of authors across different career stages. The pie chart shows the percentage of authors according to underrepresented minority status. Data in B, C and D were plotted using Prism (n=158).

### Story Readability

Readability of the stories is crucial to the objective of the journal in ensuring the stories are accessible. While the demographic analyses (Fig. 1) are based on authors (n=158), the forthcoming analysis is based on the individual stories (n=170 published by those authors). We used the industry standard Flesch reading ease score to ascertain the readability of the stories, where scores are determined by the average length of sentences (based on number of words) and the average number of syllables per word.^6^ The scale ranges from 0 to 100, with zero being very difficult to read (understandable by university graduates) and 100 being very easy to read (understandable by an average 11-year-old student). We determined that 74% of the 170 stories included in the analysis fall in the target range of 50-70, which represents readability by students in grades 8-12 (Figure 2A). When looking at the full distribution of the scores across different score bins, the 60-65 score range features the most stories (40) (Figure 2B). These results are ideal in that there are very few stories on the extremes of the score spectrum and most fall within the target readable range.

**Figure 2:**
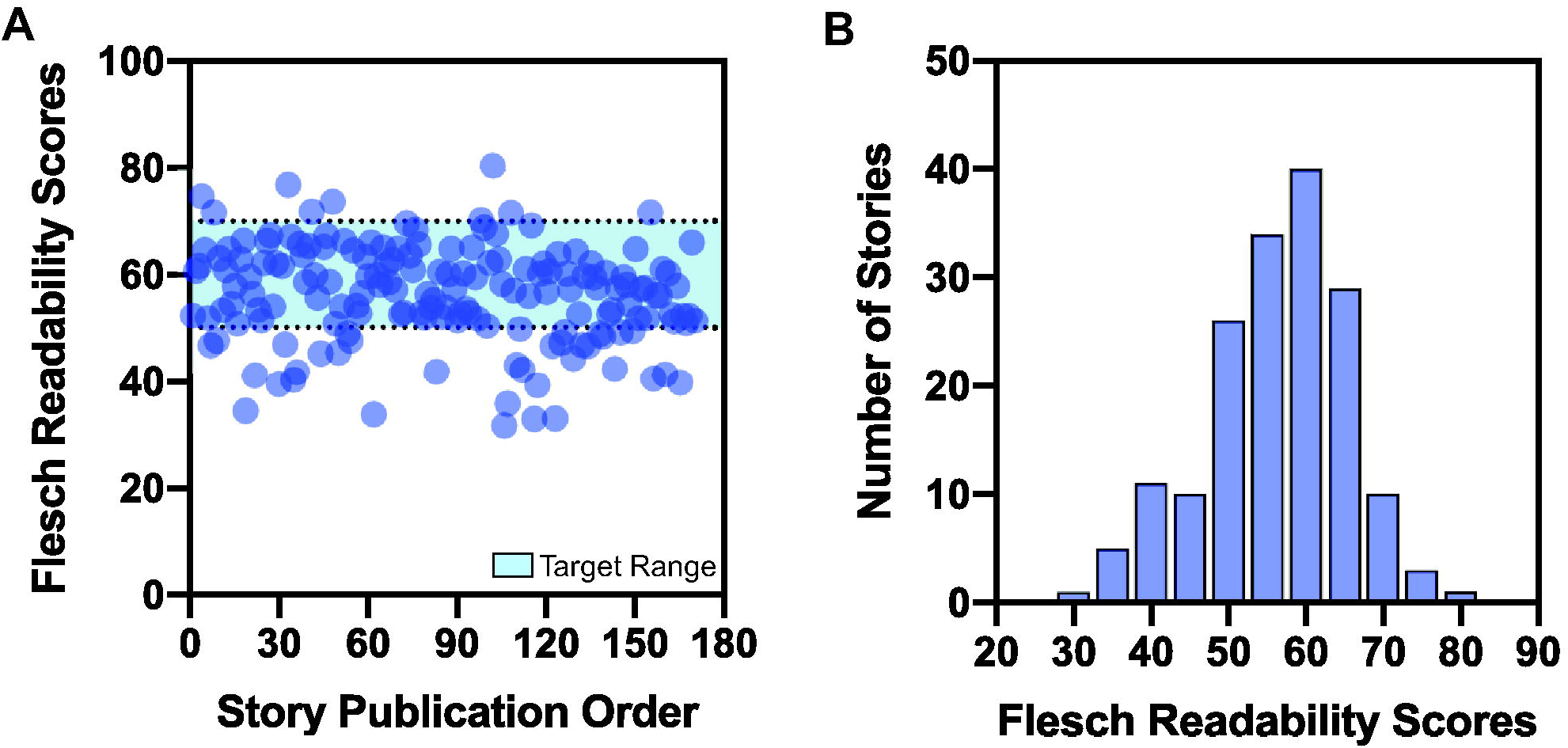
Readability of published stories. **(A)** The Flesch reading ease scores (y-axis) are plotted by publication date (x-axis). The blue shade shows the target readable range between 50-70. **(B)** Flesch reading ease scores are represented **as a histogram** (bin width = 5).

### Reader Engagement

It is important to study reader engagement to ensure that the narratives are reaching an attentive and responsive audience. To examine reader engagement, we looked at page views per story by publish date (Figure 3). We find that the majority of the stories have less than or around 500 page views, but there are a few outliers with many more views. There is a cluster of three stories around the 2500-3000 page views mark, and there is one story with almost 8000 page views. Based on the data shown in Figure 3 and qualitative analyses of the most versus least popular stories by page views, it is difficult to predict which stories will be popular. Some stories end up getting more attention than others, which can be caused by a number of variables. It is worth mentioning that a page view is counted each time the page is visited, so duplicates are included. Page views also do not factor the time spent on the page by each reader. Furthermore, a page view does not indicate whether the reader actually read the story or what impact it had on them. Nonetheless, the page view metric demonstrates a power-law distribution with a minority of stories reaching “viral” status.

**Figure 3:**
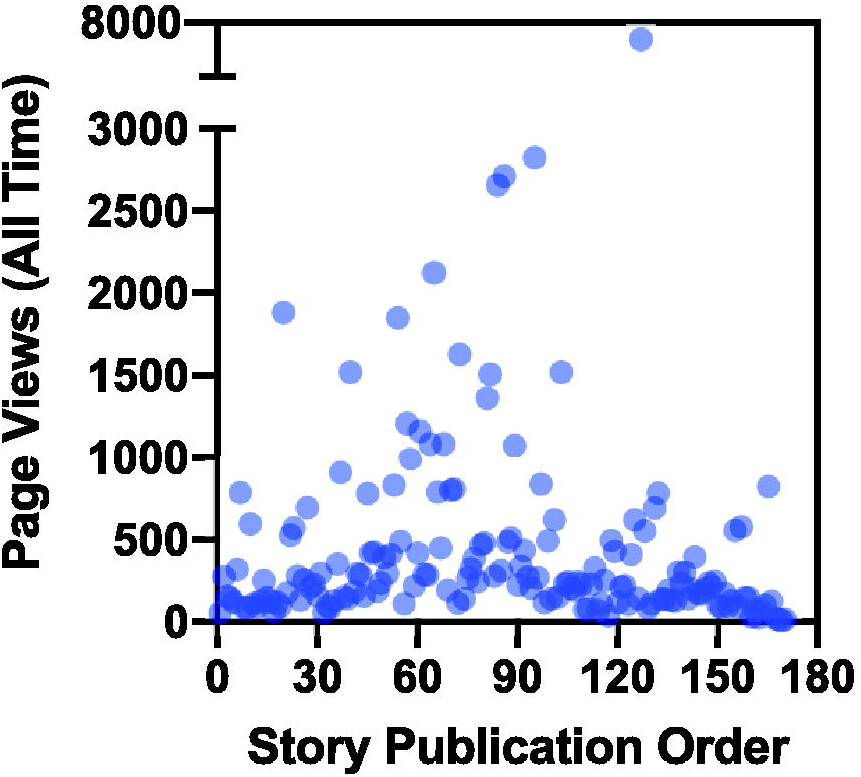
Popularity of stories as assessed by page views. Individual page views for each of the 170 stories published between December 3, 2016 and June 14, 2020 are plotted chronologically. Page view data are from Google Analytics.

### Future Directions

The analysis here provides the first deep look into one of the databases publishing diverse stories in science using a wide range of mediums. First, the demographic background analysis shows that the editorial team must continue to focus on diversifying the database of authors by actively doing outreach across various scientific fields, career stages, locations, and demographic backgrounds. This could be achieved by engaging and partnering with universities, organizations, professional societies, and also groups both within and outside the United States to increase the submission of diverse stories. Such engagement will be critical moving forward for the journal. It is also noteworthy that the majority of authors thus far have been women which could be attributed to the heightened challenges women face in STEM careers.^7^ There is still a lack of representation of women in STEM and thus it is imperative that their voices are shared broadly especially with the next generation. An important aspect of our analysis is the story readability. With over three-fourths of the stories being readable by high school students (scores over 50), it becomes possible to consider incorporating these stories into high school science curricula, especially given the known benefits of students hearing these stories.^8-10^ The stories provide perspective on what being a scientist really looks like, so focusing on exposing high schoolers to that information would be advantageous. As mentioned previously, it is tough to accurately predict the impact of a story. The page view metric we used is a blunt approach to begin to answer that question. The metric only goes so far as to provide insights into the general popularity of stories. It is worth mentioning that the average time spent by readers on each story is roughly two minutes. Again, this metric still says little about the impact and utility of the stories. With more time for analysis, it would have been useful to examine reader engagement by analyzing time spent on each story, as this would also give insights into which stories are most engaging and accessible for readers. This would also indicate which stories (if any) are the ones that receive the most thorough read compared to others.

In summary, our analysis is simply the first step and it generates a number of questions and considerations. Namely, it will be essential that other databases track and disseminate their own relevant metrics to allow both readers and other stakeholders to assess the utility of the platforms. Beginning this process could be as simple as deciding which metrics are most relevant to track for analysis. Dissemination of such data will be important for the field as the insights gained will help drive the field of storytelling in science forward. As more and more scientists tell their stories, it will be imperative that their narratives are examined. Among many critical questions is the need to come up with innovative metrics that assess the impact of storytelling in science and also answer whether the stories are increasing our understanding of what it means to be a human in science.

## Acknowledgements

We would like to thank all the members of the STEM Advocacy Institute (SAi) for their valuable feedback.

## Author Contribution Statement

R.Q.M., J.W.T., and F.J.M. jointly conceived the idea for the research project. R.Q.M. performed the initial data organization and F.J.M. verified the analytical methods used by R.Q.M. F.J.M. mentored R.Q.M. to conduct the analysis and supervised the findings of the work. All authors discussed the results and contributed to the final manuscript.

## Competing interests

None

## Data Availability

The data that support the findings of this study are available from the corresponding author upon request.

## Notes

### Competing Interest Statement

The authors have declared no competing interest.

